# Dose-dependent progression of multiple low dose streptozotocin-induced diabetes in mice

**DOI:** 10.1101/2023.04.08.536122

**Authors:** Brandon M. Bauer, Supriyo Bhattacharya, Elizabeth Bloom-Saldana, Jose M. Irimia, Patrick T. Fueger

**Affiliations:** Department of Molecular and Cellular Endocrinology, Arthur Riggs Diabetes and Metabolism Research Institute, City of Hope, Duarte, CA 91010, USA; Irell & Manella Graduate School of Biological Science, Beckman Research Institute, City of Hope, Duarte, CA, 91010, USA; Integrative Genomics Core, Beckman Research Institute, City of Hope, Duarte, CA 91010, USA; Comprehensive Metabolic Phenotyping Core, Beckman Research Institute, City of Hope, Duarte, CA 91010, USA

## Abstract

This study investigated the effects of different multiple low doses of streptozotocin (STZ), namely 35 and 55 mg/kg, on the onset and progression of diabetes in mice. Both doses are commonly used in research, and while both induced a loss of beta cell mass, they had distinct effects on whole glucose tolerance, beta cell function and gene transcription. Mice treated with 55 mg/kg became rapidly glucose intolerant, whereas those treated with 35 mg/kg had a slower onset and remained glucose tolerant for up to a week before becoming equally glucose intolerant as the 55 mg/kg group. Beta cell mass loss was similar between the two groups, but the 35 mg/kg-treated mice had improved glucose-stimulated insulin secretion in gold-standard hyperglycemic clamp studies. Transcriptomic analysis revealed that the 55 mg/kg dose caused disruptions in nearly five times as many genes as the 35 mg/kg dose in isolated pancreatic islets. Pathways that were downregulated in both doses were more downregulated in the 55 mg/kg-treated mice, while pathways that were upregulated in both doses were more upregulated in the 35 mg/kg treated mice. Moreover, we observed a differential downregulation in the 55 mg/kg-treated islets of beta cell characteristic pathways, such as exocytosis or hormone secretion. On the other hand, apoptosis was differentially upregulated in 35 mg/kg-treated islets, suggesting different transcriptional mechanisms in the onset of STZ-induced damage in the islets. This study demonstrates that the two STZ doses induce distinctly mechanistic progressions for the loss of functional beta cell mass.

## INTRODUCTION

Type 1 diabetes mellitus (T1D) results from the autoimmune-mediated destruction of the insulin-producing beta cells in the Islets of Langerhans in the pancreas. Without a cure for T1D, the field continues to rely on experimental model systems to identify potential therapeutics. Two of the main model systems employed are the non-obese diabetic mouse and beta cell destruction models using chemical agents such as alloxan and streptozotocin (STZ).

STZ is an alkylating agent that is selectively cytotoxic to beta cells (1-3). It is a common experimental tool to study the loss of beta cell mass in rodents as well as beta cell replacement strategies (4). STZ induces DNA damage, oxidative stress, and apoptosis, leading to chronic pancreatic islet inflammation, insulitis, and insulin deficiency, which resembles features of human T1D. However, the complete mechanisms governing beta cell mass loss in STZ-induced models of diabetes remain incompletely understood (2). Whereas a single injection of a high dose (≥150 mg/kg) of STZ can induce a near-complete necrotic ablation of beta cells within 24 hours and overt hyperglycemia within 48 hours, multiple administrations of a lower dose of STZ results in a more gradual loss of beta cell mass and delayed hyperglycemia, a phenotype not directly attributable to the drug cytotoxicity (4). Therefore, multiple low dose administration of STZ is often preferred as a model of human disease since it induces beta cell changes that model diabetes progression such as immune infiltration and beta cell dysfunction (5, 6). Additionally, low-dose STZ has been used to study beta cell regeneration and repair (7-9). However, there remains little uniformity in the field when it comes to selecting a low STZ dose.

The original work to introduce the multiple dose technique used 40 mg/kg (10), but doses as low as 30 mg/kg or as high as 55 mg/kg are commonly used. We sought to compare the impact of a multiple low dose (55 mg/kg) to a multiple very low dose (35 mg/kg) administration of STZ. Our goal was to better understand how the STZ concentration alters the metabolic, morphologic, functional, and transcriptomic progression of STZ-induced rodent beta cell dysfunction as a model of T1D.

We identified key differences in disease progression between a 35 and 55 mg/kg multiple low dose of STZ. Whereas 55 mg/kg-treated animals became glucose intolerant within three days following STZ administration, 35 mg/kg-treated animals remained glucose tolerant for several days longer than 55 mg/kg-treated animals despite an equivalent loss of beta cell mass. Using hyperglycemic clamps, the gold standard for quantifying insulin secretion *in vivo*, we confirmed that 35 mg/kg-treated mice partially retained glucose-stimulated insulin secretion for an extended period following treatment compared to 55 mg/kg-treated mice. There were also significant differences in gene expression between the two treatment groups, including dose-dependent shifts in differentially expressed genes and pathways as well as a subset of genes altered in opposite directions. This work clarifies the impact of STZ concentration and can help researchers to select an appropriate dose for their experiments to properly model disease progression and improve intervention models to retain or restore beta cell mass in diabetes.

## METHODS

### Animal Studies

All animals were maintained in accordance with City of Hope Institutional Animal Care and Use Committee protocols (#16047) in accordance with the Guide for the Care and Use of Laboratory Animals (11). Only male C57BL/6J were used due to the varying effectiveness of STZ in female mice. All mice were housed in individually ventilated cages with 3-5 mice/cage. Animals were maintained on 12-h light:dark cycles with *ad libitum* access to water and standard rodent chow diet.

STZ (Sigma-Aldrich S0130-1G; St. Louis, MO, USA) was administered for five consecutive days by intraperitoneal (IP) injections of STZ dissolved right before the injection in saline at either 35 or 55 mg/kg body weight. Equivalent volumes of saline were used in control animals. Glucose tolerance tests were performed on 10–12-week-old male mice following a five-h fast; glucose (1.5 g/kg) was delivered by IP injections. Blood glucose measurements were made using an AlphaTRAK 2 (Abbott Laboratories; Abbott Park, IL, USA) glucometer from a tail nick blood sample.

### Quantification of Beta Cell Mass

Pancreata were removed from mice following CO_2_ asphyxiation, and tissues were fixed in 10% formalin overnight. Pancreas slices were processed into formalin blocks with assistance from the Solid Tumor Pathology Core at City of Hope, and 6 µm section slides were made from the paraffin blocks. Paraffin removal was performed by immersion in xylene, and slide rehydration was achieved by serial washes in graded ethanol solutions. Antigen retrieval was performed by boiling slides in pH 6.0 citrate buffer (H3300; Vector Laboratories; Newark, CA, USA) for 10 min. Slides were subsequently allowed to cool to room temperature for 30 minutes. Slides were blocked with serum free protein block (Dako X0909; Agilent Technologies; Santa Clara, CA, USA). Insulin was labeled with primary guinea pig anti-insulin antibody (1:500; ab195956; Abcam; Waltham, MA, USA) at 4C overnight and detected with secondary biotinylated goat anti guinea pig antibody (Vector Laboratories; BA7000, 1:500) followed by Vectastain Elite ABC HRP detection kit (Vector Laboratories; PK-6100) and DAB (K346811-2; Agilent Technologies). Slides were counter stained with Mayer’s Hematoxylin (TA-125-MH; Epredia; Kalamazoo, MI, USA).

Slides were imaged using a NanoZoomer 2.0HT (Hamamatsu Photonics; Shizuoka, Japan) slide scanner and analyzed using QuPath (V.0.3.2). The pixel identification feature of QuPath (12, 13) was used to automatically identify and quantify either total pancreas area or insulin positive area in a blinded fashion. All images were checked for the presence of positive identification of artifact and were manually corrected when necessary.

### Hyperglycemic Clamps

Surgical implantation of carotid artery and jugular vein catheters and hyperglycemic clamp procedures to assess *in vivo* beta cell function were performed by the Comprehensive Metabolic Phenotyping Core at City of Hope according to techniques previously described (14-16). Briefly, mice were anesthetized with isoflurane and surgery was conducted using aseptic techniques. An arterial catheter was inserted through an incision, terminating approximately 9.5 mm into the carotid artery lumen, and was secured by proximal and distal ligatures. A second catheter was inserted into the right jugular vein terminating approximately 10 mm into the vessels lumen and secured with proximal and distal ligatures. The free ends of each catheter was tunneled subcutaneously toward the back of the mouse, exteriorized through a small incision in the interscapular region, and connected to the mouse antenna for sampling access (MASA; made in-house) device. The MASA device was then secured with suture, making blood sampling/infusion ports easily accessible from the interscapular region of the mouse. Animals were provided analgesia and fluids and monitored post-operatively.

After four days of recovery, mice that returned to within 10% of pre-surgical body weight were attached to a two-way swivel (Instech, Plymouth Meeting, PA), allowing animals to remain conscious and freely moving throughout the procedure. Animals received a primed (50 mg/kg/min for 2 min) followed by variable infusion of 50% dextrose and continuous infusion of saline-washed erythrocytes throughout the experiment, to achieve hyperglycemia (target: 270 mg/dl, or 18 mM) and to maintain a constant hematocrit, respectively. Arterial blood glucose was monitored during the clamp every 10 min (∼5 µl whole blood). Blood samples (∼100 µl each) were collected at −15, −5, 5, 10, 15, 30, 60, 90 and 120 minutes to measure blood insulin (Sensitive Rat Insulin Kit, #SRI-13K; MilliporeSigma; Burlington, MA, USA) following manufacturer instructions. Mice were euthanized at the conclusion of the study, and the pancreas collected for histology.

### Islet Isolation

Islets were isolated using methods previously described by Stull *et al*. (17) with freshly prepared CIzyme RI collagenase and protease solution (VitaCyte 005-1030; Indianapolis, IN, USA). After isolation, healthy islets were manually picked and cultured overnight at 37° with 5% CO_2_ in RPMI supplemented with 10% FBS, 5 mM HEPES (Fisher Scientific, BP310-500), and penicillin/streptomycin (50 units/ml and 50 μg/ml, respectively; Gibco/ThermoFisher Scientific, 15-140-122; Waltham, MA, USA).

### RNA Sequencing

Approximately 75-100 healthy islets per mouse were picked by hand, washed in cold PBS, and then resuspended in 350 ul of RLT with 1%BME. RNA was isolated using RNeasy micro (Qiagen 7404; Hilden, Germany) kits according to manufacture instructions.

RNA sequencing libraries were prepared with Kapa RNA mRNA HyperPrep kit (Kapa Biosystems, Cat KR1352; Cape Town, South Africa) according to the manufacturer’s protocol. The final libraries were validated with the Agilent Bioanalyzer DNA High Sensitivity Kit and v1.5 with the paired end mode of 2x101cycle. Real-time analysis (RTA) v3.4.4 software was used for base calling.

### Statistical Analysis

All data are presented as means ± SEM. RNA data were normalized to control conditions and presented as relative expression. Adjusted area-under-the-curve (AUC) was calculated using the trapezoid method. Student’s *t*-test (unpaired, two-tailed unless otherwise stated) or ANOVA (with Bonferroni post hoc tests) were performed using GraphPad Prism software (La Jolla, CA, USA) to detect statistical differences. *p < 0.05* was considered statistically significant.

## RESULTS

### Divergent progression of metabolic dysfunction is STZ dose dependent

To better define the impact of streptozotocin dose on the progression of functional beta cell mass loss in mice, ten-week-old male mice were administered intraperitoneal injections of STZ for five consecutive days at either a low dose of 55 mg/kg or a very low dose of 35 mg/kg. Before the STZ administration (control) and on the third, tenth, and twentieth days following the final dose of STZ, glucose tolerance tests were performed by challenging 5-hour fasted mice with 1.5 g/kg glucose by IP injection (**Figure 1A**).

**Figure 1:**
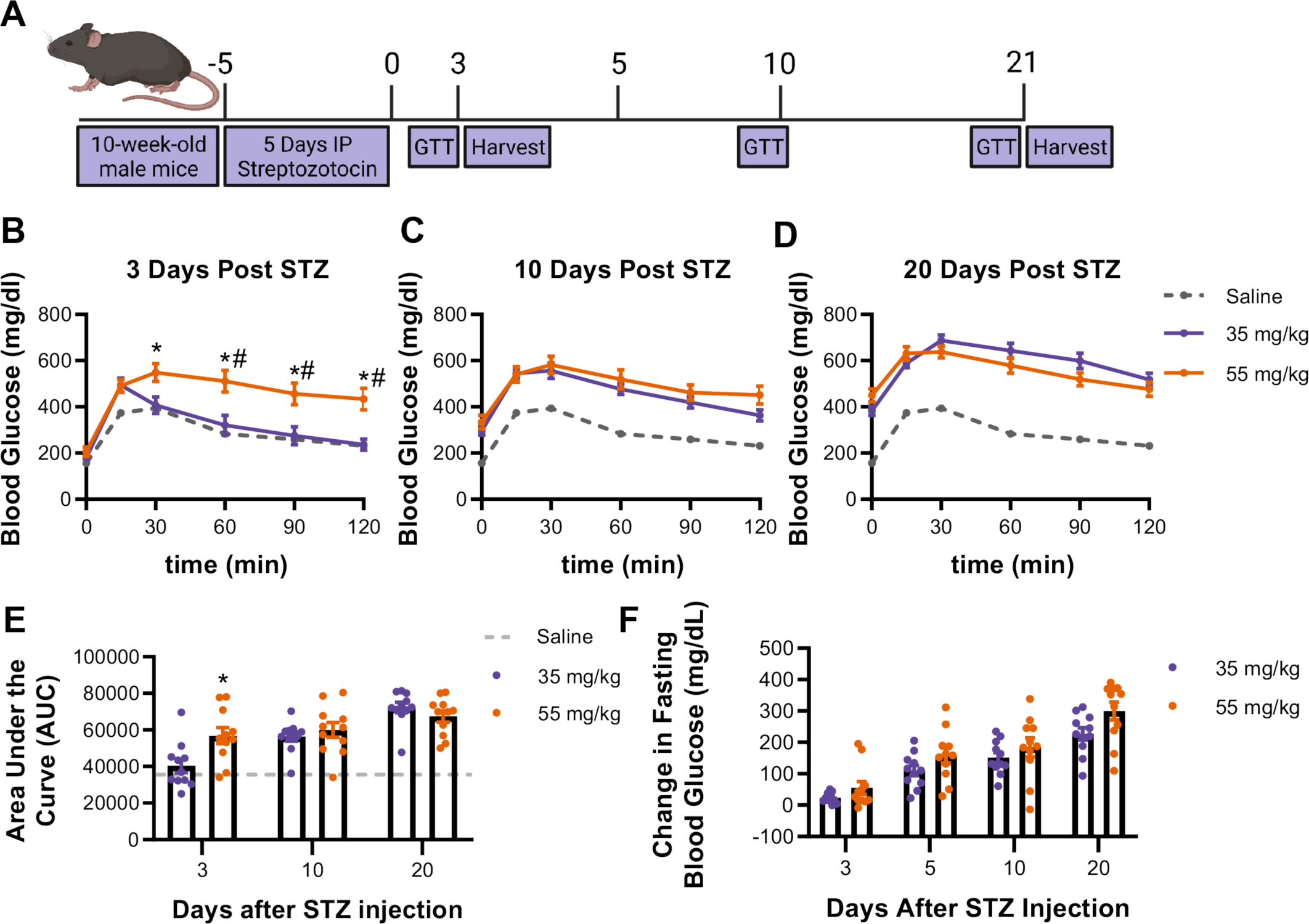
Impact of 35 mg/kg vs 55 mg/kg dose STZ on glucose tolerance and fasting blood glucose. **A.** Mice were treated for 5 consecutive days with either 35 mg/kg or 55 mg/kg STZ. **B.** Glucose tolerance was assessed by an IP-GTT (1.5 g/kg glucose) at 3 days following the final day of STZ. Additional GTTs were performed at **C.** 10 days and **D.** 20 days following the final day of STZ. **E.** The area under the curve (AUC) of the GTTs was found at 3 days and similar at 10- and 20-days post STZ. **F.** Fasting blood glucose on days 3, 5, 10, 20 (2-way ANOVA, *p<0.05 vs control, #p<0.05 vs 35 mg/kg STZ, n=6-7 mice, mean ± SEM)

At three days, post STZ there were significant differences in glucose tolerance between the 35 and 55 mg/kg-treated mice (**Figure 1B**). Whereas the 35 mg/kg-treated mice resembled control (i.e., saline-treated) animals, 55 mg/kg-treated mice were significantly less glucose tolerant than both control and 35 mg/kg-treated mice. The difference in glucose tolerance, intolerant by the tenth day following treatment, and through twenty days following STZ administration (**Figure 1C-E**). Five-hour fasting blood glucose continued to increase throughout the three weeks following STZ administration in both STZ treatment groups (**Figure 1F**). There remained a slight, albeit not statistically significant, trend in decreased fasting blood glucose between the very low and low dose STZ-treated animals.

### Beta cell mass loss progresses similarly under both STZ regimens

To establish the impact of STZ dose on beta cell mass we quantified pancreatic beta cell area via immunohistochemical labeling of insulin-positive cells (**Figure 2A**). By the third day following STZ beta cell area had already declined by greater than 40% in both the 35 and 55 mg/kg-treated mice (**Figure 2B-D**). Interestingly, after the initial loss of beta cell mass within the first several days after STZ treatment, beta cell area did not continue to decline at the same rate despite the progressive worsening in glucose tolerance. Both beta cell area (**Figure 2C-D**) and mass (**Figure 2E**) remained similar between the two treatment groups, which suggests loss of beta cell mass, as judged by imaging of insulin-positive cells, alone was not the primary factor impacting disease progression after the initial insult induced by STZ. We posited there might instead be differences in beta cell function, rather than beta cell mass, between the two treatment groups.

**Figure 2:**
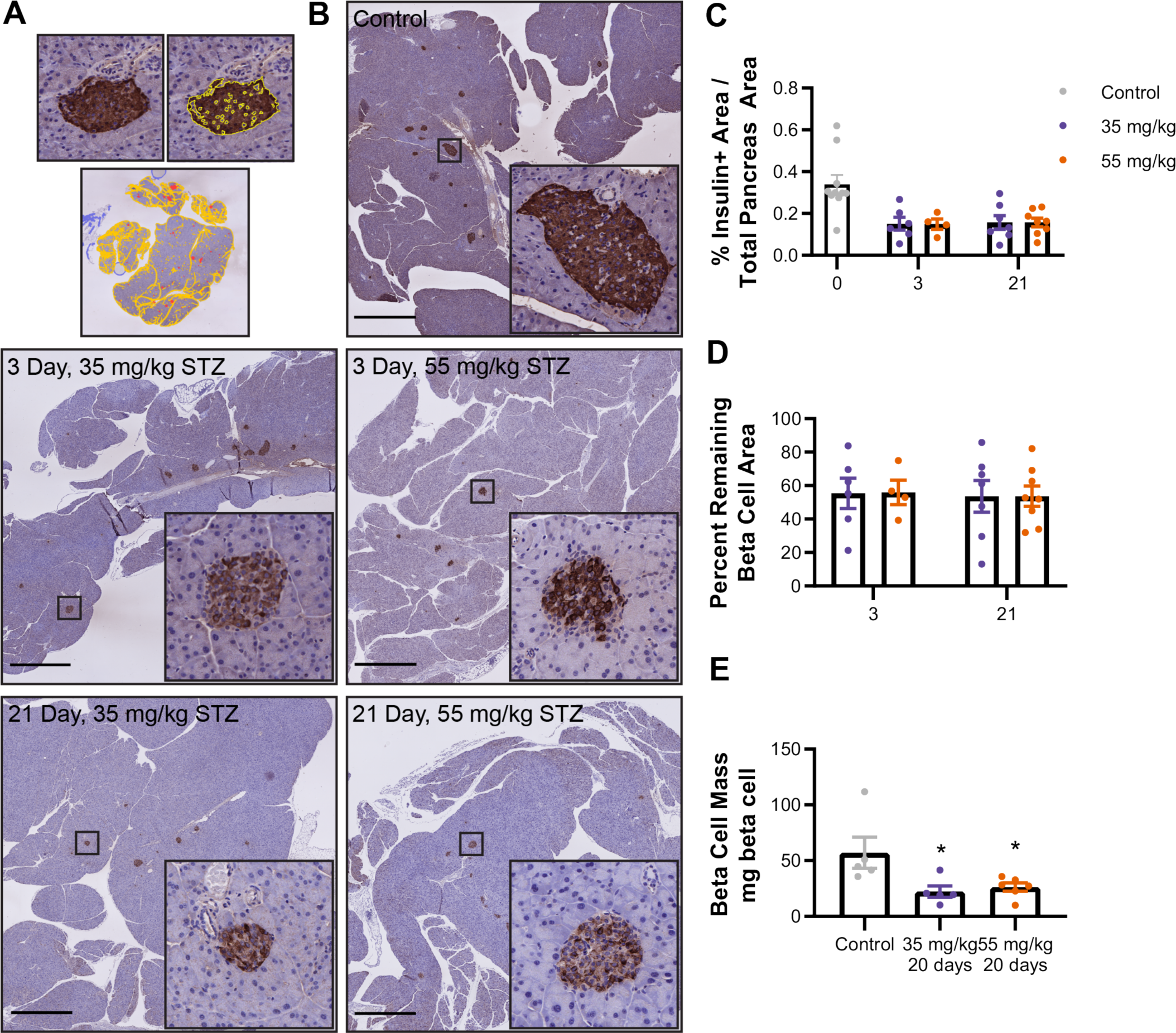
STZ-induced loss of Beta Cell Mass is comparable between 35 mg/kg and 55 mg/kg STZ. **A.** Model of insulin positive area detection by QuPath. **B.** Representative images of Pancreas sections from STZ-treated mice labeled for insulin by IHC and counterstained with hematoxylin. Shown are whole pancreas images (scale bar = 1mm) or individual islets (20x magnification). Pixel identification in QuPath was used to identify whole pancreas area and insulin positive tissue area. **C.** There was a significant decrease in insulin + area after STZ without any differences between the two doses. **D.** Overall there was about a 40% decrease in beta cell mass following treatment with STZ for both doses **E.** Beta cell mass (beta cell area x pancreas weight) was decreased compared to controls but similar between STZ doses. (2-way ANOVA, *p<0.05 vs control, #p<0.05 vs 35 mg/kg STZ, n=6-7 mice, mean ± SEM)

### Hyperglycemic clamps elucidate dose-dependent loss of insulin secretion

Given the improved glucose tolerance at 3 days post-STZ in the 35 mg/kg compared to the 55 mg/kg group, despite similar remaining beta cell area, we hypothesized that the 35 mg/kg-treated mice retained beta cell function in the remaining beta cells. To determine how conscious animals, considered the gold standard methodology for quantifying *in vivo* insulin secretion (18). We, therefore, treated mice with streptozotocin, or saline as control, for five days as previously described. On the fifth day of injection (day 0 of post-treatment), we surgically implanted catheters, and after allowing the animals to recover from surgery, hyperglycemic clamp experiments were performed on the fourth day after surgery recovery to coincide with the period of preserved glucose tolerance we observed in the 35 mg/kg-treated mice (**Figure 3A**).

**Figure 3:**
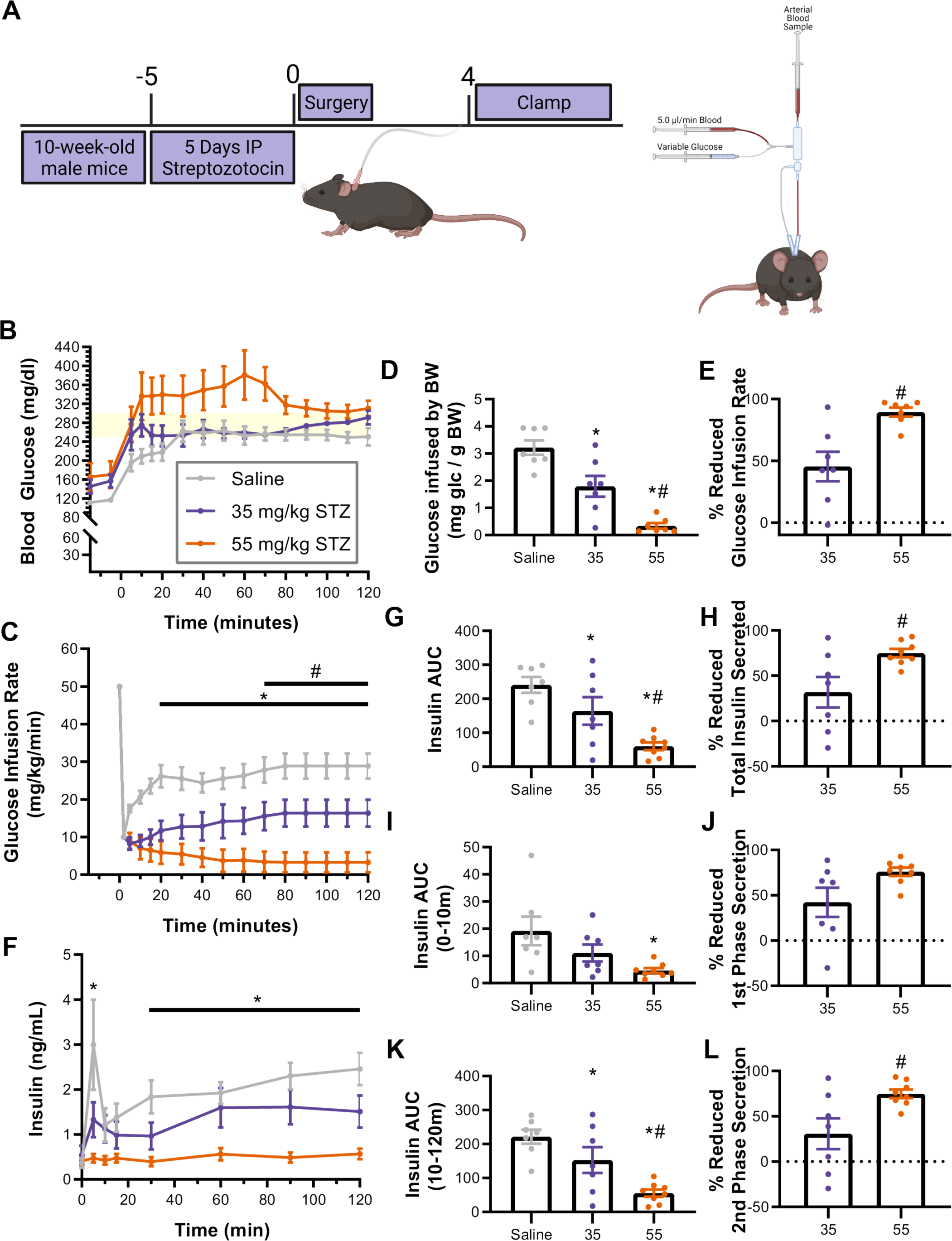
35 mg/kg STZ has prolonged period of glucose stimulated insulin secretion before complete loss of beta cell function. **A.** Timeline demonstrating the administration of STZ followed by surgical implantation of catheters and hyperglycemic clamp procedure. **B.** Mice were maintained at a constant hyperglycemic target between 250 and 300 mg/kg for 2 hours. **C.** Glucose was infused at a variable rate (mg/kg/min) to maintain hyperglycemia. **D.** Overall glucose infused was calculated as the Area Under the Curve of the GIR graph and **E.** total glucose infused (mg glucose / kg body weight). **F.** Insulin secretion, normalized to pre-glucose-infused levels, were found at 5, 10, 15, 30, 60, 90, and 120 minutes. **G.** Area under the curve of insulin secretion was significantly lower in 55 mg/kg STZ than CON or 35 mg/kg STZ. **H.** During first phase (0-10 minutes) and **I.** second phase (10-120 minutes) AUC was significantly in 35 but not 55 mg/kg STZ. (2-way ANOVA, *p<0.05 vs saline, #p<0.05 vs 35 mg/kg STZ, n=6-7 mice, mean ± SEM)

The goal of the clamp is to achieve a stable blood glucose within the target range (250-300 mg/dL), reaching steady-state in glucose infusion rate (GIR). Coinciding with results reported in (**Figure 1**), the STZ-treated animals from both groups had higher basal/fasting blood glucose compared to the control animals, suggesting the surgical procedure did not interfere with the STZ-induced glucose intolerance. All animals achieved a steady state by 10 minutes at which point they maintained stable blood glucose throughout the procedure (**Figure 3B**).

GIR was decreased in both STZ-treated groups compared to the saline-treated control group after 10 minutes through the end of the clamp. In the mice treated with 55 mg/kg STZ GIR values were also decreased compared to the 35 mg/kg group after 50 minutes (**Figure 3C**). Overall, total glucose infused was significantly lower in 35 mg/kg treated animals than saline-treated animals, and both parameters were even lower in 55 mg/kg treated mice (**Figure 3D**). The 35 mg/kg treated mice had 45.4±11.9% overall reduction in GIR and the 55 mg/kg treated mice had 89.5±3.7% reduction (**Figure 3E**).

Saline-treated animals secreted the highest levels of insulin throughout the procedure (**Figure 3F**). Whereas-35 mg/kg treated animals tended to secrete less insulin, the only individual timepoint that was significantly lower was at 2 min. Overall insulin AUC (**Figure 3G**) was different between all groups; however, whereas 55 mg/kg-treated mice had a 75.0±13% loss of insulin, 35 mg/kg-treated had only a 33.8% loss (**Figure 3H**).

We also analyzed the loss of insulin AUC in the first phase (0-10 min, **Figure 3I-J**) versus the second phase (10-120 min, **Figure 3K-L**) of insulin release. In 35 mg/kg-treated animals, whereas the peak insulin secretion at 2 min was lower than saline treated controls, first phase insulin AUC was not significantly lower than saline treated animals (**Figure 3I**). On the other hand, in 55 mg/kg-treated animals insulin levels were not increased from baseline and were significantly lower than saline and 35 mg/kg groups throughout most of the clamp period (**Figure 3F-L**). Moreover, in 55 mg/kg animals first phase insulin release was nearly completely abolished (**Figure 3I-J**) and second phase was robustly reduced (**Figure 3K-L**). Overall, these data suggest that glucose-stimulated insulin secretion was at least partially preserved at 4 days in 35 mg/kg treated animals, and 55 mg/kg treated animals had a more rapid loss of both first and second phase insulin secretion compared to 35 mg/kg STZ-treated animals.

### Transcriptomic changes associated with STZ dosage

Given that the 55 and 35 mg/kg STZ-treated mice had differences in beta cell function shortly after STZ, we sought to characterize transcriptional changes that might be driving these differences in beta cell function. We isolated primary mouse islets on the third day following treatment with STZ or saline, allowed the islets 12 hours to recover from the isolation process, and then performed bulk RNA sequencing. Whereas there were differences in the genes that were disrupted, the majority of differentially expressed genes in 35 mg/kg-treated mice were similarly differentially expressed in 55 mg/kg-treated animals. 55 mg/kg-treated mice however had substantially more differentially expressed genes. We identified 596 upregulated and 460 downregulated genes in the 35 mg/kg-treated animals. In 55 mg/kg-treated animals there were nearly 5x more differentially expressed genes with 2384 genes upregulated and 2304 downregulated (**Figure 4A-B**).

**Figure 4:**
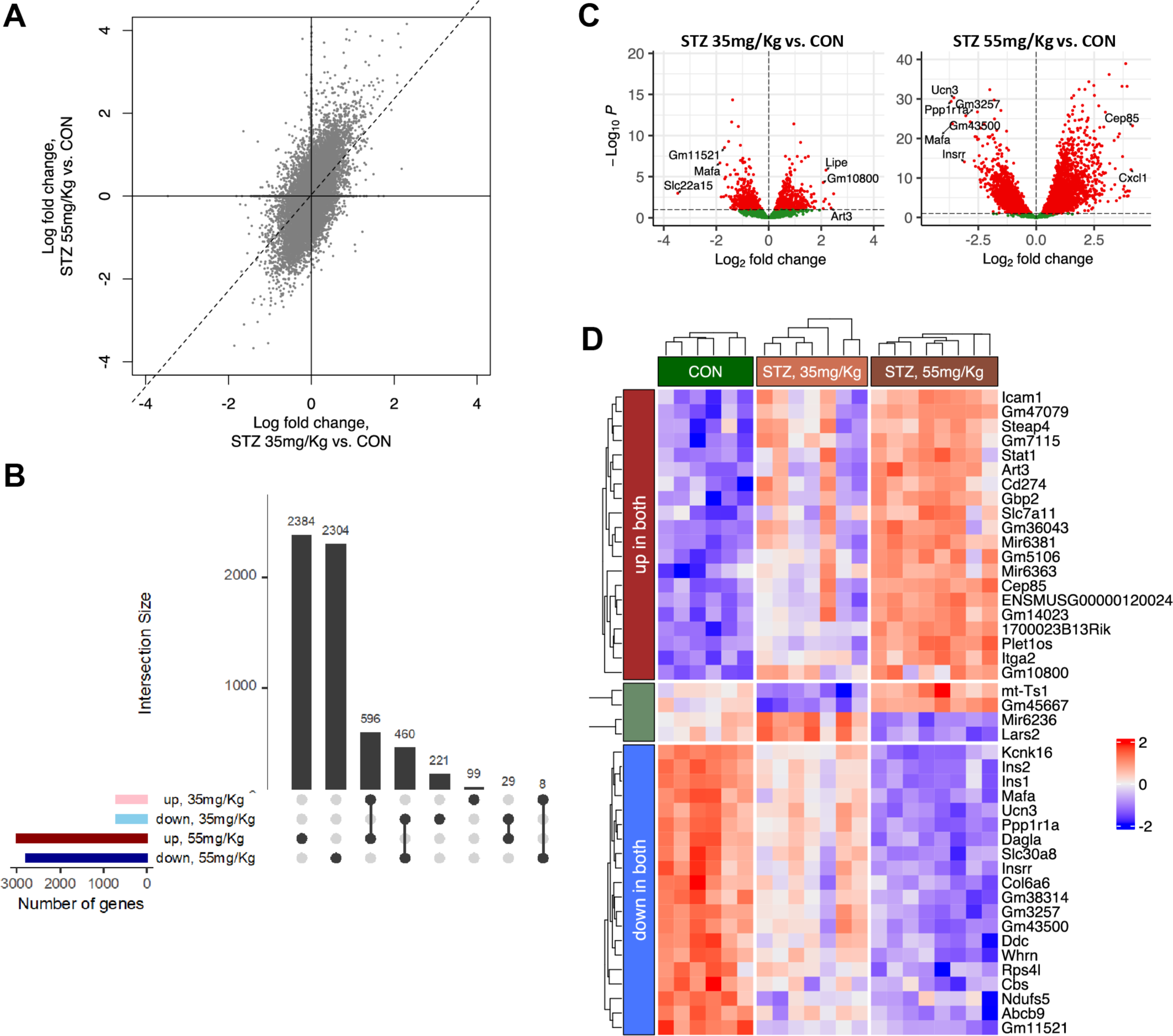
Impact of STZ dose on gene expression. **A.** Correlation among log fold changes of gene expressions between the two STZ doses. Dotted line represents the linear regression fit, along with the corresponding r2 highlighted in the plot. **B.** Upset plot showing the overlap of up and downregulated genes (adjusted pvalue < 0.1) between the two STZ groups. **C.** Volcano plots of expression log fold changes vs statistical significance for the two STZ groups. The top up and down genes are labeled in each plot. **D.** Heatmap highlighting the expressions of genes that show similar and opposite direction of change between the two STZ groups. Each column represents one mouse sample. Genes are first filtered by statistical significance (adjusted pvalue < 0.1) and those showing log fold change > ±1 are retained. Top 10 genes in up/down category are shown, while only four genes were found to change expression in the opposite direction between the 35 and 55 mg/Kg STZ groups. Each row is individually scaled between ±2.

Similar to other studies examining the impact of STZ on gene expression (19), we identified increased expression of genes associated with cytokines response and inflammation (e.g., Gbp2/3, Stat1, Cxcl1) and extracellular matrix remodeling (e.g., Itga2); key genes related to beta cell identity or insulin secretion (e.g., Ucn3, MafA, Ins1) and metabolic regulation (e.g., Fh1) were decreased (**Figure 4 C-D**). Interestingly, genes and pathways that were upregulated tended to be more upregulated in 35 vs 55 mg/kg group, and the genes that were downregulated were more downregulated in 55 vs 35 mg/kg group, suggesting a dose-dependent effect of STZ on gene expression. There was a small subset of genes that were altered in opposing directions with the different STZ doses; 29 genes were down in 35 mg/kg and up in 55 mg/kg and 8 genes that were up in 35 mg/kg and down in 55 mg/kg (**Figure 4B,D**).

### STZ dose-dependent differential expression of genes and pathways related to endocrine cell function and survival

Despite the overall similarities in differentially expressed genes (**Figure 4**), the 35 mg/kg and 55 mg/kg animals had distinct patterns of gene expression. In both groups, there were similar patterns of differentially expressed genes involved in pathways related to metabolism/ATP production/respiration, insulin/hormone secretion, and immune/stress response. Notably there were dose dependent differences in the degree to which these pathways were disrupted (**Figure 5A-B**).

**Figure 5:**
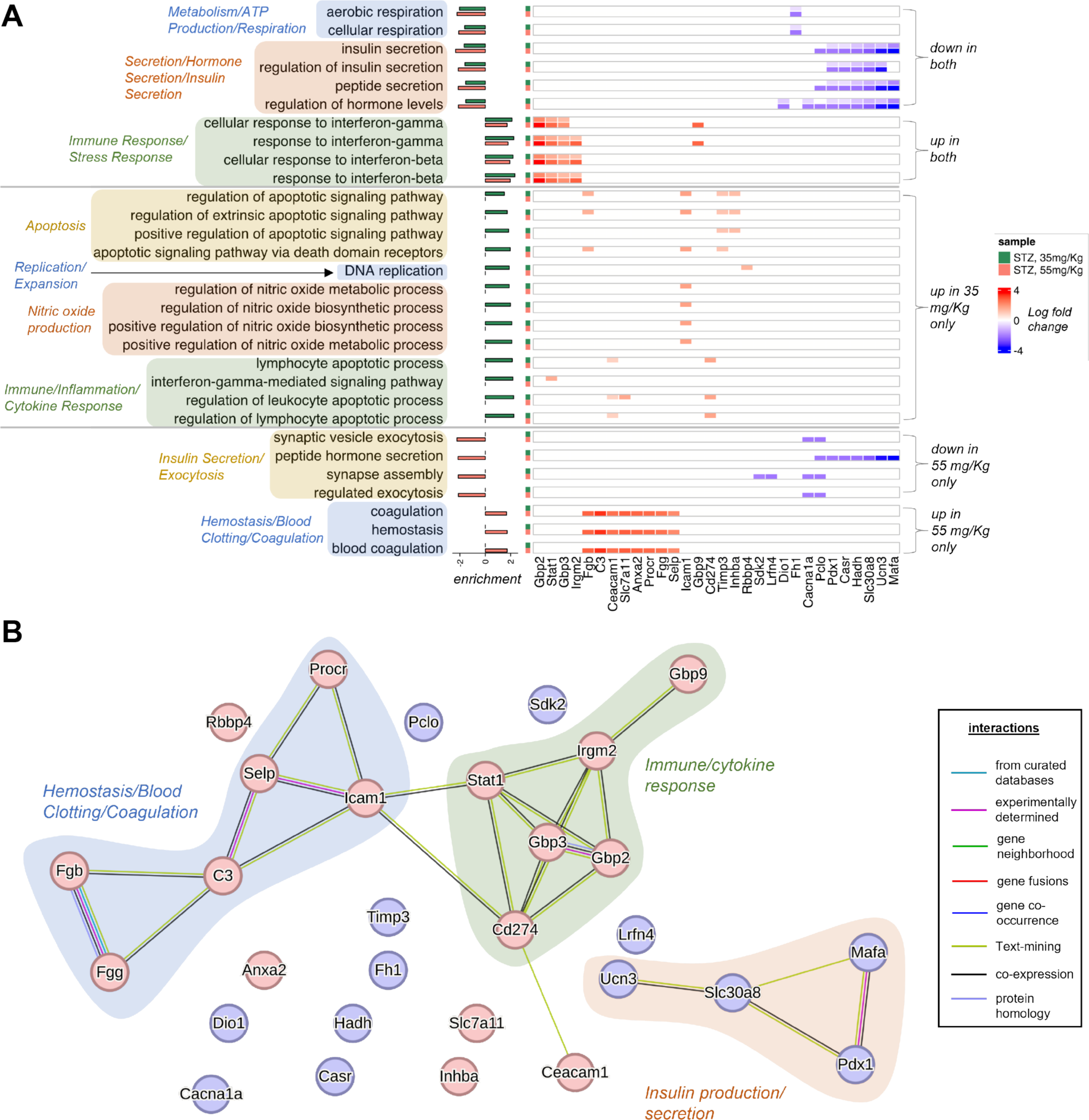
Pathway analysis of 35 mg/kg STZ vs 55 mg/kg STZ. **A.** Heatmap of top differentially expressed genes (|log fold change| > 1.5) among the enriched biological processes in STZ, 35 and 55 mg/Kg samples, grouped process type. For each process, the log fold changes in both samples are shown, color coded on the left according to sample type (35 mg/Kg: green, 55 mg/Kg: red). Barplot on the left reflects normalized enrichment scores (NES) as obtained from geneset enrichment analysis. **B.** Protein interaction network from STRING database involving the genes highlighted in panel A. Subnetworks related to major enriched processes are highlighted in different shades. Up and down genes are colored red and blue respectively.

We characterized the pathways between 35 and 55 mg/kg-treated animals that were differentially expressed and found that compared to the 55 mg/kg-treated animals, 35 mg/kg-treated animals had increased expression of genes related to pathways involving apoptosis, replication/expansion, nitric oxide production, and inflammatory/cytokine response. Alternatively, 55 mg/kg had down regulation of insulin exocytosis pathways not observed in 35 mg/kg as well as increased expression of pathways related to hemostasis/blood clotting/coagulation (**Figure 5A**). Changes in these expression of genes in these pathways support evidence of a greater loss of insulin secretion in 55 mg/kg-treated mice compared to 35 mg/kg-treated mice observed in hyperglycemic clamps (**Figure 3**).

## DISCUSSION

Rodent models of diabetes such as STZ-induced diabetes are essential tools to examine the progression of functional beta cell mass loss in diabetes and are commonly used to evaluate strategies to prevent or reverse beta cell loss (4). Administration of multiple low doses of STZ highlights many features of human T1D progression such as immune infiltration, beta cell dysfunction, regeneration, and repair (5-9); this experimental model allows researchers to evaluate strategies to prevent or reverse beta cell loss. Despite the widespread use of multiple low-dose STZ-induced models of diabetes, there remains little uniformity in the dose used.

Additionally, there are still uncertainties regarding the mechanisms driving disease progression in STZ-induced models of diabetes (2, 3). We demonstrate here that STZ dose is a critically important factor to the onset and progression of STZ-induced diabetes models. We identified significant differences in disease progression between 35 and 55 mg/kg STZ doses. Given the widespread issues with reproducibility of animal models in scientific research (20-22), researchers should, therefore, be aware of the impact of a chosen STZ dose and carefully select and report the dose used.

We discovered that despite similar levels of beta cell loss, 35 mg/kg-treated animals remained glucose tolerant for significantly longer than 55 mg/kg-treated littermates. We posited the 35 mg/kg-treated animals may have a longer period of sustained beta cell function. To test this hypothesis, we performed hyperglycemic clamps in STZ-treated mice, which we believe is the first time such experiments have been reported. Data supported our hypothesis and demonstrated that 4 days following STZ administration, 35 mg/kg-treated mice secreted significantly more insulin than 55 mg/kg-treated, and therefore, were able to better manage hyperglycemia despite having similar levels of beta cell mass loss. Thus, the two doses allow one to uncouple beta cell loss from function.

From the hyperglycemic clamps, we were able to distinguish between first and second phases of insulin secretion in both 35 and 55 mg/kg STZ-treated groups. This aspect of our study is critical given that loss of first-phase insulin secretion, which is defined as the insulin secreted in the first 10 minutes following an increase in blood glucose, is an important hallmark of diabetes onset and represents a clinically important indicator of beta cell dysfunction (23). We determined that whereas there was a decrease in peak insulin secretion in 35 mg/kg treated mice, the total AUC for 35 mg/kg mice was not significantly lower than saline controls. Both 35 and 55 mg/kg STZ-treated mice did, however, have a loss of second phase insulin secretion compared to saline-treated control mice. Although mechanistically distinct from the first phase, second phase insulin secretion is physiologically significant in that it can be sustained for hours after hyperglycemia is induced, accounts for the majority of insulin secretion following a meal and is a major regulator of glucose homeostasis. Additional work should be done to fully describe the mechanisms impacting both first and second phase insulin secretion following multiple low dose STZ.

To elucidate the molecular mechanisms governing disease progression in the STZ-treated mice we assessed the impact of STZ dose on gene expression from isolated islets. We concluded the changes in gene expression were generally similar between the doses and aligned with previously published transcriptomic reports (19, 24). However, we discovered significant dose-dependent changes in gene expression that had not previously been reported. The higher dose of STZ induced changes in the expression of nearly 5x more genes than 35 mg/kg STZ. Additionally, within the similarly disrupted pathways, we found that the 55 mg/kg STZ-treated mice tended to have a greater downregulation of transcriptionally suppressed pathways whereas 35 mg/kg STZ-treated animals tended to have a greater upregulation of induced pathways. Comparing the doses revealed that several key pathways were in fact differentially expressed. 35 mg/kg-treated animals had relative increased expression of genes in pathways regulating replication/expansion, nitric oxide production, apoptosis, and inflammatory/cytokine response. Conversely, 55 mg/kg-treated animals had decreased expression of insulin secretion/exocytosis related genes.

We acknowledge that both the 35 and 55 mg/kg doses can be valuable tools for researchers. A 55 mg/kg dose reliably induces a substantial but incomplete loss of functional beta cell mass to induce glucose intolerance in a relatively short period (less than 3 days). Alternatively, a lower dose of 35 mg/kg provides investigators with an extended period to study beta cell dysfunction and may therefore be a more appropriate model to test interventions intended to preserve or reverse functional beta cell mass loss, especially for those strategies requiring some time to develop. In addition, the 35 mg/kg dose may represent, at least in mice, a time period analogous to T1D diagnosis; on the other hand, the 55 mg/kg dose would align more closely with the weeks and months after diagnosis. Given the recent success of the first approval for a treatment that prolongs C-peptide levels in newly diagnosed people with T1D (25, 26), and presumably more to come (27, 28), more well-defined preclinical models can facilitate efforts to develop more early intervention therapies.

We do note that our findings have several limitations resulting from the animals we studied. All experiments were restricted to male C57BL/6J mice that were around 10 weeks old. Male mice are preferred to female mice, which are known to be largely resistant to STZ-induced diabetes and have significant differences in how they respond to STZ (29-31). Likewise, different strains of mice have been documented to respond to STZ in significantly different ways with some requiring significantly larger doses to induce diabetes (29, 32-34). In addition, beta cell dysfunction (35), as well as the efficiency of STZ-induced beta cell damage, is known to be age-dependent (36). However, these findings can be a stepping-stone to understanding the importance of dose and future studies can, and already have begun to, elucidate sex and strain-specific differences in the response to STZ.

We conclude that variations in multiple low dose STZ regimens can elicit significant differences in the progression of functional BCM loss in mice. Although a range of doses are used to induced diabetes, dose selection has significant impacts on the rate at which beta cell function declines. Whereas beta cell mass progression appeared to decline similarly between the 35 and 55 m/kg doses, the loss of beta cell function progressed significantly faster in 55 mg/kg treated mice compared to 35 mg/kg treated mice. The higher dose of STZ also elicited a more robust changes in gene expression. The prolonged development of diabetes with the lower dose of STZ could be advantageous to testing the efficacy of emerging therapies in pre-clinical models. Thus, researchers should take careful precautions in selecting, preparing, and reporting STZ doses when using STZ-induced models of diabetes.

## Author contributions itemized list

Conceptualization, BMB and PTF; methodology, BMB, JI, SB, and PTF; formal analysis, BMB, SB, EB-S, JI, and PTF; investigation, BMB, EB-S, JI, and PTF; resources, PTF; data curation, BMB and SB; writing - original draft preparation, BMB and PTF; writing - review and editing, BMB, SB, EB-S, JI, and PTF; visualization, BMB and SB; supervision, PTF; project administration, BMB and PTF.; funding acquisition, PTF. All authors have read and agreed to the published version of the manuscript.

## ACKNOWLEDGELMENTS

This research was funded by National Institutes of Health grant DK099311 to PTF and Innovation Awards from the Wanek Family Project to Cure Type 1 Diabetes to PTF. This work utilized services from the Center for Comparative Medicine and Solid Tumor Pathology, Integrated Genomics, and Light Microscopy Cores at the City of Hope, which were supported by the National Cancer Institute of the National Institutes of Health grant P30CA033572.

